# Structure of *Allium* lachrymatory factor synthase elucidates catalysis on sulfenic acid substrate

**DOI:** 10.1101/142687

**Authors:** Takatoshi Arakawa, Yuta Sato, Jumpei Takabe, Noriya Masamura, Masahiro Kato, Morihiro Aoyagi, Takahiro Kamoi, Nobuaki Tsuge, Shinsuke Imai, Shinya Fushinobu

## Abstract

Natural lachrymatory effects are invoked by small volatile *S*-oxide compounds. They are produced through alkene sulfenic acids by the action of lachrymatory factor synthase (LFS). Here we present the crystal structures of onion LFS (*Ac*LFS) revealed in solute-free and two solute-stabilized forms. Each structure adopts a single seven-stranded helix-grip fold possessing an internal pocket. Mutagenesis analysis localized the active site to a layer near the bottom of the pocket, which is adjacent to the deduced key residues Arg71, Glu88, and Tyr114. Solute molecules visible on the active site have suggested that *Ac*LFS accepts various small alcohol compounds as well as its natural substrate, and they inhibit this substrate according to their chemistry. Structural homologs have been found in the SRPBCC superfamily, and comparison of the active sites has demonstrated that the electrostatic potential unique to *Ac*LFS could work in capturing the substrate in its specific state. Finally, we propose a rational catalytic mechanism based on intramolecular proton shuttling in which the microenvironment of *Ac*LFS can bypass the canonical [1,4]-sigmatropic rearrangement principle established by microwave studies. Beyond revealing how *Ac*LFS generates the lachrymatory compound, this study provides insights into the molecular machinery dealing with highly labile organosulfur species.

**Significance statement:** Crushing of onion liberates a volatile compound, *syn*-propanethial *S*-oxide (PTSO), which causes lachrymatory effect on humans. We present the crystal structures of onion LFS (*Ac*LFS), the enzyme responsible for natural production of PTSO. *Ac*LFS features a barrel-like fold, and mutagenic and inhibitory analyses revealed that the key residues are present in the central pocket, harboring highly concentrated aromatic residues plus a dyad motif. The architecture of *Ac*LFS is widespread among proteins with various biological functions, such as abscisic acid receptors and polyketide cyclases, and comparisons with these homologs indicate that unique steric and electronic properties maintain the pocket as a reaction compartment. We propose the molecular mechanism behind PTSO generation and shed light on biological decomposition of short-lived sulfur species.

## Introduction

Organosulfur compounds are often accumulated in vegetables, and they have improved the quality of human life in the form of particular sensory experiences as well as nutritional and health benefits (1–5). Pungent smells evoked by the *Allium* plants stem from cytosolic (+)-*S*-alk(en)yl cysteine sulfoxides (CSOs), which are produced abundantly from γ-glutamyl-*S*-alk(en)yl cysteines (6). CSOs are odorless precursors, but disruption stresses such as injury, biting and cutting allow them to contact vacuolar C-S lyase (alliinase; EC 4.4.1.4), after which spontaneous decomposition and polymerization occur (Fig. 1). The population of mostly volatile end products [*e.g*., dialk(en)yl thiosulfinates (**5**), cepathiolanes (**6**), cepaenes (**7**), polysulfides (**8**), and zwieberanes (**9**)] contributes to determining alliaceous smells and flavors (7, 8).

**Fig. 1.**
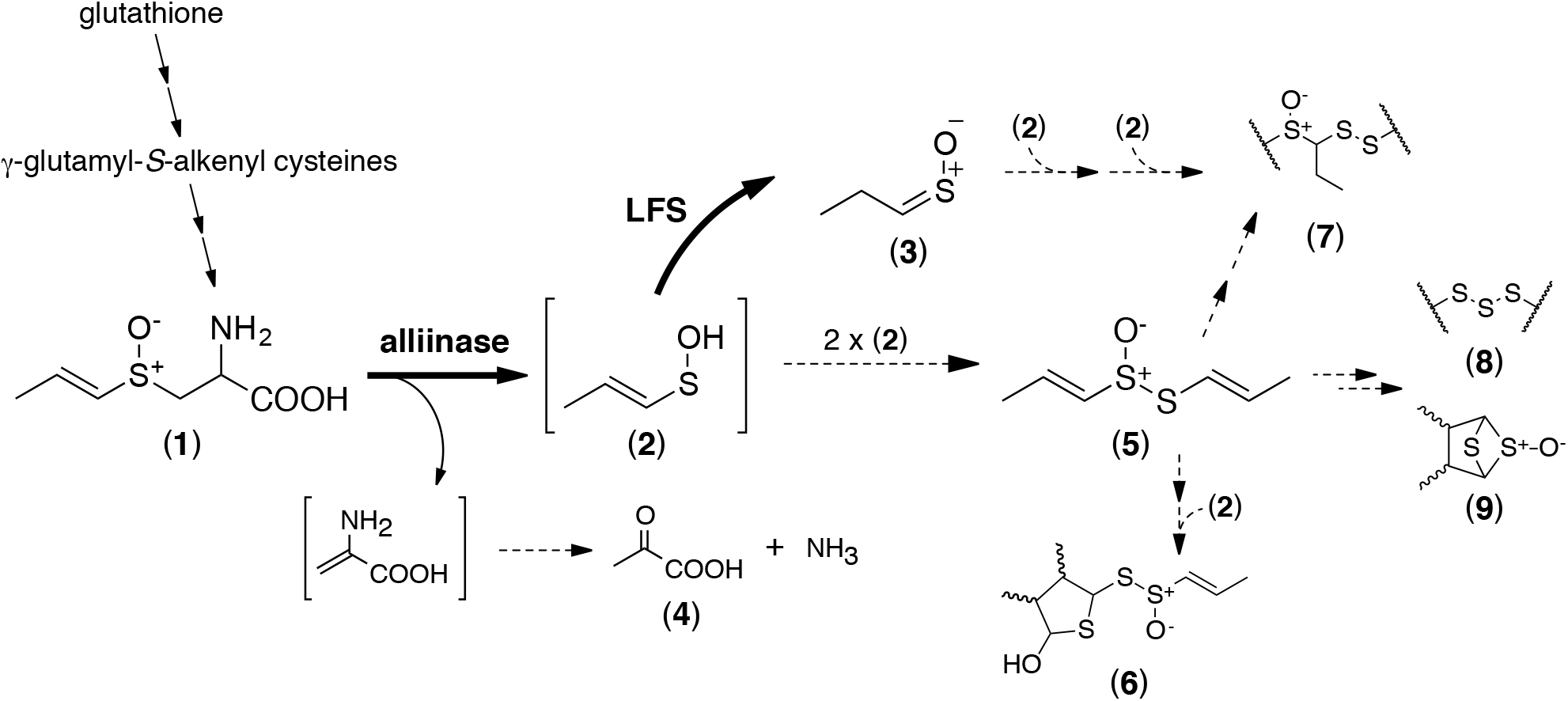
Chemical utilization pathways of γ-glutamyl-*S*-alkenyl cysteines via *trans*-1-PRENCSO in crushed onion. Solid and dashed arrows represent enzymatic and non-enzymatic processes, respectively. Brackets indicate undetectable materials. Numbered chemical names are noted in the main text, except for pyruvic acid (**4**).

In addition, crushing of bulb onion (*Allium cepa*) and other limited plants induce weeping. *syn*-Propanethial *S*-oxide (PTSO) (**3**) is a prototype of lachrymatory agents (9) and generated by the conjunct catalysis initiated by the alliinase-mediated cleavage of *trans*-*S*-1-propenyl l-cysteine sulfoxide (*trans*-1-PRENCSO/isoalliin) (**1**) (7). Lachrymatory factor synthase (LFS; EC 5.3.-.-) serves on the subsequent conversion to PTSO, and thus branches the common spontaneous degradation pathway of CSOs (10) (Fig. 1). Onion LFS (*Ac*LFS, 169 aa) is assumed to be an irreversible isomerase against (*E*)-1-propene-1-sulfenic acid (1-PSA) (**2**), but molecular machinery of *Ac*LFS remains unveiled, only implicating that the two residues (R71 and E88) associate in this catalysis (11). Before the discovery of LFS, Block and colleagues have established a canonical theory of PTSO formation through 1-PSA, termed [1,4]-sigmatropic rearrangement (Fig. 5*A*), employing flash vacuum pyrolysis (FVP) (12). This non-enzymatic, intramolecular pericyclic reaction-based principle proceeds under vacuum and heat, while *Ac*LFS mediates an equivalent reaction under ambient conditions and without the assistance of any cofactors.

LFS is a unique enzyme because it acts on a the short-lived intermediate. Sulfenic acid (R-SOH) is one of the reactive sulfur species (RSS), which is completely sensitive to thiol and sulfenyl groups as well as oxidants due to its bifacial preference as a nucleophile and an electrophile (13, 14). Sulfenic acids appear as a protein modification of cysteine residues (15–17) and function in specific regulated activities (18–24). However, they have rarely been identified as substrates due to them being too labile to be detected, with an estimated half-life of several milliseconds (25) and not more than a minute for methane sulfenic acid generated in the vacuum gas phase (26). In organisms lacking LFS genes, for example, garlic (*Allium sativa*), free sulfenic acids are condensed immediately, and thus are just destined to produce dialk(en)yl thiosulfinates (2, 6). Gene silencing of onion LFS derives increases of (**5**) and (**9**) isomers, displaying a metabolite composition similar to that of garlic (8, 27).

Here, we present the crystal structures of *Ac*LFS. They enable us to elucidate the natural mechanism by which coincident and spontaneous RSS reactions are selectively modified by this enzyme.

## Results

### Overall structure and confirmation of the active site

Crystal structures of *Ac*LFS were determined at a resolution of 1.7 to 2.1 Å (Table S1). In the present models, the N-terminal region (1–22), which is not required for the activity (11), was not determined due to conformational disorder. *Ac*LFS (23–169) formed a single globular domain with dimensions of 40×40×43 Å, consisting of a seven-stranded antiparallel β-sheet (S1–S7), a long α-helix (H3), two flanking helices (H1, H2), and connecting loops (Fig. 2*A*). The molecular architecture in which the b-sheet bends so as to wrap H3 is known as a helix-grip fold (28).

**Fig. 2.**
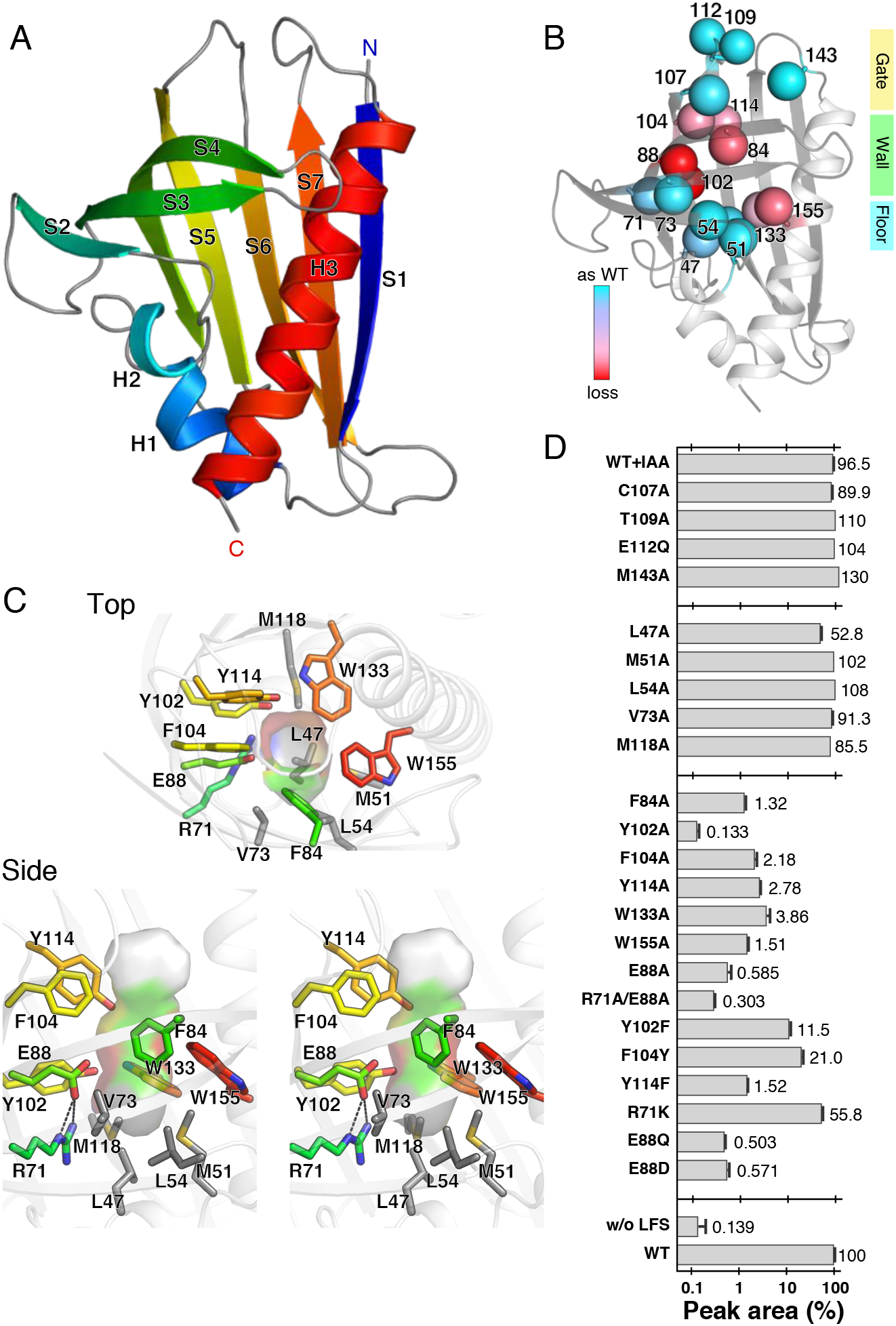
Structure and the active site of *Ac*LFS. (*A*) Overall structure displayed by schematic representation in spectral colors. (*B*) Positions of mutated residues colored by impacts on the LFS activity. Spheres are γ-atoms colored in cyan (none) to red (significant), based on the activities of respective alanine variants, R71K and E112Q. (*C*) Pocket-lining residues. Views from the entrance (top) and the direction of the parallel to longitudinal axis (bottom, stereoview). The side chains of Wall and Floor layers (sticks) are shown with the cartoon drawing of the main chain. The carbon atoms of Wall are colored as in (*A*) and those of Floor are in gray. (*D*) List of variants and relative productivities. Error bars represent standard deviations of at least triplicate measurements.

There are five cysteine residues on the molecular periphery of *Ac*LFS (Fig. S1*A*). Inactivation of the free thiols by an excess amount of iodoacetamide (IAA) caused no significant change in PTSO generation (Fig. 2*D*). Hence, the catalysis does not involve any cysteine residues, which is also supported by the result that the activity of C107A remained at the same level as that of intact *Ac*LFS.

*Ac*LFS possesses an elongated pocket extending 18 Å from the molecular surface. This pocket can be divided into three layers, termed Gate, Wall, and Floor (Fig. 2*B*). Wall and Floor are pocket-lining apparatuses provided by side chains of S1, S3–7, and H2–3, and consist mainly of aromatic and aliphatic residues, respectively (Fig. 2*C*, S4). A series of mutagenesis analyses proved that the active site resides in the Wall region. Specifically, variants of Wall residues (F84A, Y102A, F104A, Y114A, W133A, W155A) exhibited equivalent reductions in activities to less than 10%, while those of Floor (L47A, M51A, L54A, V73A, M118A) and a part of Gate (C107A, T109A, W112Q, M143A) maintained the activities roughly at the level of the wild type (Fig. 2*D*). Bulky side chains of the Wall residues appear to be necessary for the catalysis since substitutions that lead to maintenance of the aromatic rings (*e.g*., Y102F and F104Y) partially retained the activities compared with their alanine mutants.

R71 and E88, which are suspected of being involved in catalysis (11), were positioned proximally at the boundary of Wall and Floor. E88A and E88Q comparably caused severe decreases in the activities to less than 1% of that of the wild type. We could not measure the activity of R71A due to protein instability. Instead, a double mutant, R71A/E88A, was examined and showed that the activity declined more than that of E88A. Although R71 influenced the activity, it was unlikely to contact 1-PSA directly, since it is located slightly away from the pocket surface (Fig. 2*C*, 3*A*). It should be noted that E88D showed a low level of activity like E88A, and R71K maintained activity at nearly half that of the wild type (Fig. 2*D*). These findings indicate that the catalysis is ruled strictly by the geometry of the two residues. The charge groups of these residues were associated tightly with two hydrogen bonds (Fig. 2*C*). Closely aligned acid and base residues can form a dyad motif in which a proton is localized between the residues, and induce a decrease in p*K*_a_ of the acid residue (29).

**Fig. 3.**
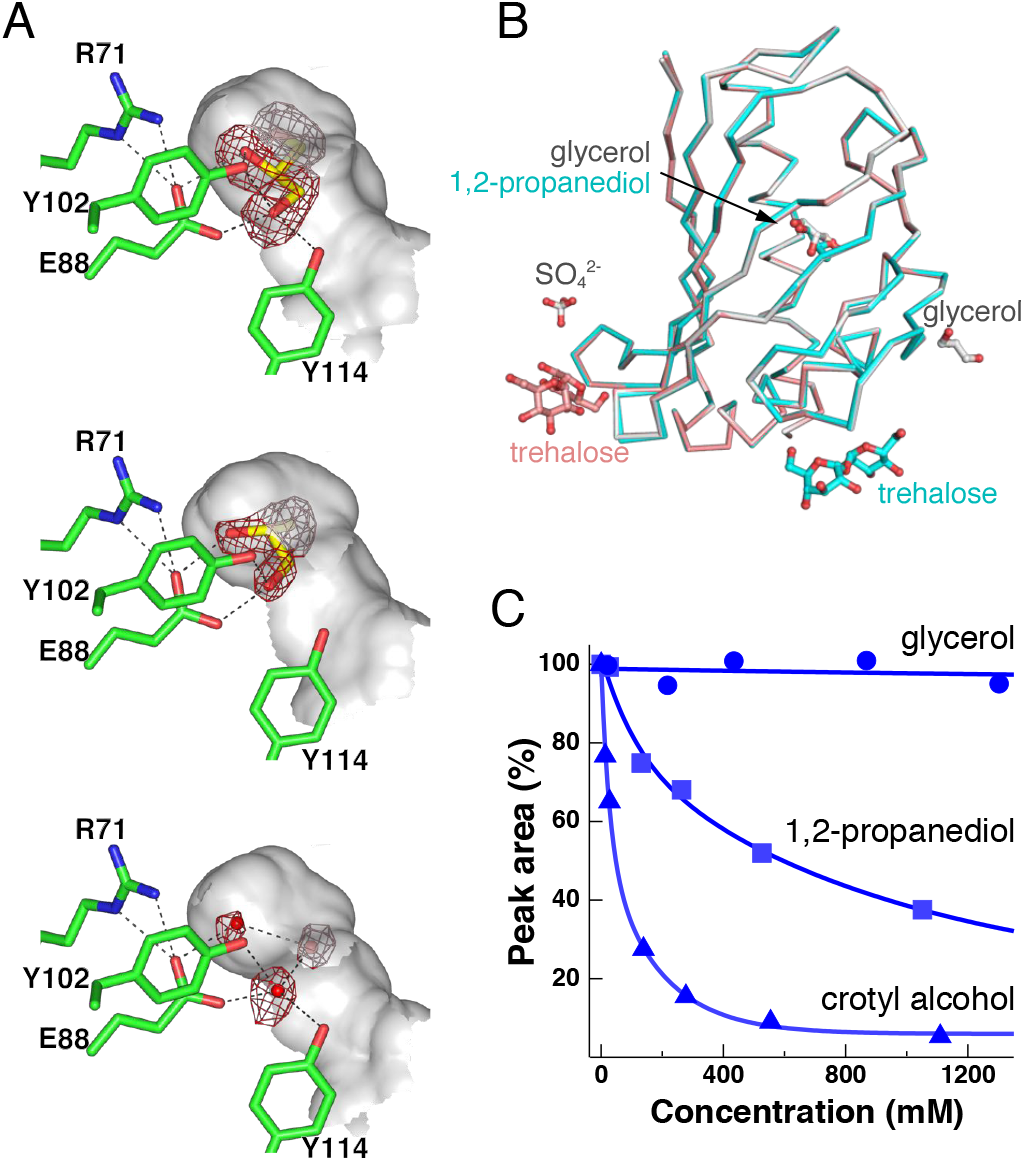
Solute molecules inside the pocket. (*A*) Non-protein electron densities found at the active site. *m*|*F*_o_|–*D*|*F*_c_| omit maps (red, 3σ) are displayed. Surfaces colored in gray are provided by the hydrophobic residues. Atoms suggested to form hydrogen bonds are connected with dashes. (*B*) Superposition of a protomer of each model. Traces of Cα atoms are taken from the data of +glycerol (white), +1,2-propanediol (pink), and solute-free (cyan). Sticks represent exogenous compounds observed on the crystals. (*C*) Inhibition curves of *Ac*LFS in the presence of glycerol, 1,2-propanediol, and crotyl alcohol.

The hydroxyl group of Y114 is an additionally important factor, as suggested by the activities of Y114A and Y114F being at the same level as those of the alanine mutants of Wall residues. In turn, the activities of Y102F and F104Y were restored compared with those of the alanine mutants (Fig. 2*D*). In the condensed hydrophobic side chains, Y114, Y102, R71, and E88 were arrayed to form a polar zone, in which Y114 is positioned near the entrance. Therefore, its hydroxyl group is the first contacting point for solute molecules (Fig. 3A). From these findings, we concluded that Y114 and E88, supported by R71, are the primal key residues for the catalysis of *Ac*LFS, and the surrounding aromatic residues work in auxiliary regulation of this activity.

### Alcohols as inhibitors

In this study, three *Ac*LFS structures were determined together with exogenous molecules (Table S1). The first one is glycerol, which originates from cryoprotectant solution (Fig. 3*A*, top). A glycerol molecule is stabilized near the bottom of the pocket, and two of the three hydroxyl groups are fixed by hydrogen bonds to E88, Y102, and Y114. The carbon moiety orients toward W133 and W155; thus, the conformation is also maintained by hydrophobic interactions. When trehalose was used as a cryoprotectant instead of glycerol, three water molecules were observed at the corresponding site of the glycerol (Fig. 3*A*, low). Subsequently, 1,2-propanediol was added to the cryoprotectant solution to obtain its complex structure (Fig. 3*A*, middle). A 1,2-propanediol molecule was observed in the pocket and adopts a conformation like that for glycerol, except for the lack of a hydrogen bond to Y114. Trehalose molecules were seen only at minor peripheral sites; thus, the pocket is unlikely to accommodate the disaccharide molecules (Fig. 3*B*).

Next, the inhibition of *Ac*LFS by these compounds was examined. 1,2-Propanediol and crotyl alcohol exhibited concentration-dependent decreases of PTSO production (Fig. 3*C*). They are deduced to behave as competitive inhibitors, considering that 1,2-propanediol was stabilized at the active site. We further investigated the inhibition patterns of cognate compounds, and demonstrated that the efficiencies are determined by the number of hydroxyl groups (Fig. S2); while linear alcohols (C_*n*_OH_2*n*+2/2*n*_; *n* ≤ 6) inhibited the reaction with half-inhibition concentration (IC_50_) values from 20 to 80 mM, diol compounds [ethylene glycol (**19**), 1,2-propanediol (**20**), and 2-methyl-2,4-pentanediol (**21**)] did so at least at 600 mM. Glycerol (**22**) did not affect the activity, even at excess concentrations (Fig. 3*C*). These findings indicate that the active site of *Ac*LFS prefers alcohols rather than glycerol and diols, and suggest that alcohols stay in the pocket with disordered conformations due to perturbation of their aliphatic portions. Linear alcohols with larger aliphatic chains were prone to having lower IC_50_ values, except for 1-pentanol (**12**) and *cis*-pentene-1-ol, (**17**) (Fig. S2*A*). Concerning an exchange of solute molecules with 1-PSA at the active site, the flexible linear alcohols would gain less entropy than diols and glycerol, which would gain entropy due to the release from the two or more hydrogen bonds (Fig. 3*A*). Assuming that the differential entropies are the major factor behind the net free-energy changes upon replacements, the diols and glycerol are feasibly stable, but poor inhibitors of *Ac*LFS.

## Discussion

### Comparison with structural homologs

The seven-stranded helix-grip fold is an architectural feature shared by members of the recently established superfamily, START/RHO alpha C/PITP/Bet v 1/CoxG/CalC (SRPBCC; cl14643) (30). LFS presents no significant sequence homologies with other proteins, but a structure similarity search highlighted 3 out of 23 SRPBCC subsets (Table S2, Fig. S3*A*). Pyrabactin-resistant protein-like proteins (PYLs) [maximal score for *Arabidopsis thaliana* PYL10 (31)] are intracellular receptors for a plant hormone, abscisic acid (ABA). Ginseng major latex-like protein 151 (MLP151), which is thought to be associated with lysophosphatidic acid regulation, is a member of the PR-10/Bet v 1-type plant allergens (32, 33). ZhuI of *Streptomyces* sp. R1128 (34) belongs to the aromatase/cyclase components of bacterial type II polyketide synthases (PKCs) and catalyzes the C7–C12-specific first ring-closing reaction toward octaketide–acyl carrier protein conjugate (35).

Each SRPBCC protein has a central cavity that provides a platform for hydrophobic molecules, as observed in complex structures of PYLs and PKCs (36, 37). Although no potent ligand of MLP151 has been identified, extra non-protein electron densities were found inside the pocket in the MLP151-cognate allergen proteins (38, 39). Compared with PYL10 and ZhuI, *Ac*LFS has a small cavity (Fig. 4*D*). These differences are apparently brought about by displacements of L1, S3, and S4 (Fig. S3*B*). In PYL proteins, L1 corresponds to the Pro-cap region, which arises from a conformational change upon the binding of ABA and constructs part of the binding site for downstream regulator proteins (36, 40). On the other hand, the Cα traces of *Ac*LFS were identical among the present solute molecule complexes; thus, no conformational changes were indicated during catalysis (Fig. 3*B*). The displacement seems not only to ensure the narrow entrance of *Ac*LFS, in which bulky hydrophobic residues gather on the Gate layer (Fig. S3*C*), but also to restrict the dimensions of their original ligands and substrates (Fig. S3*A*).

**Fig. 4.**
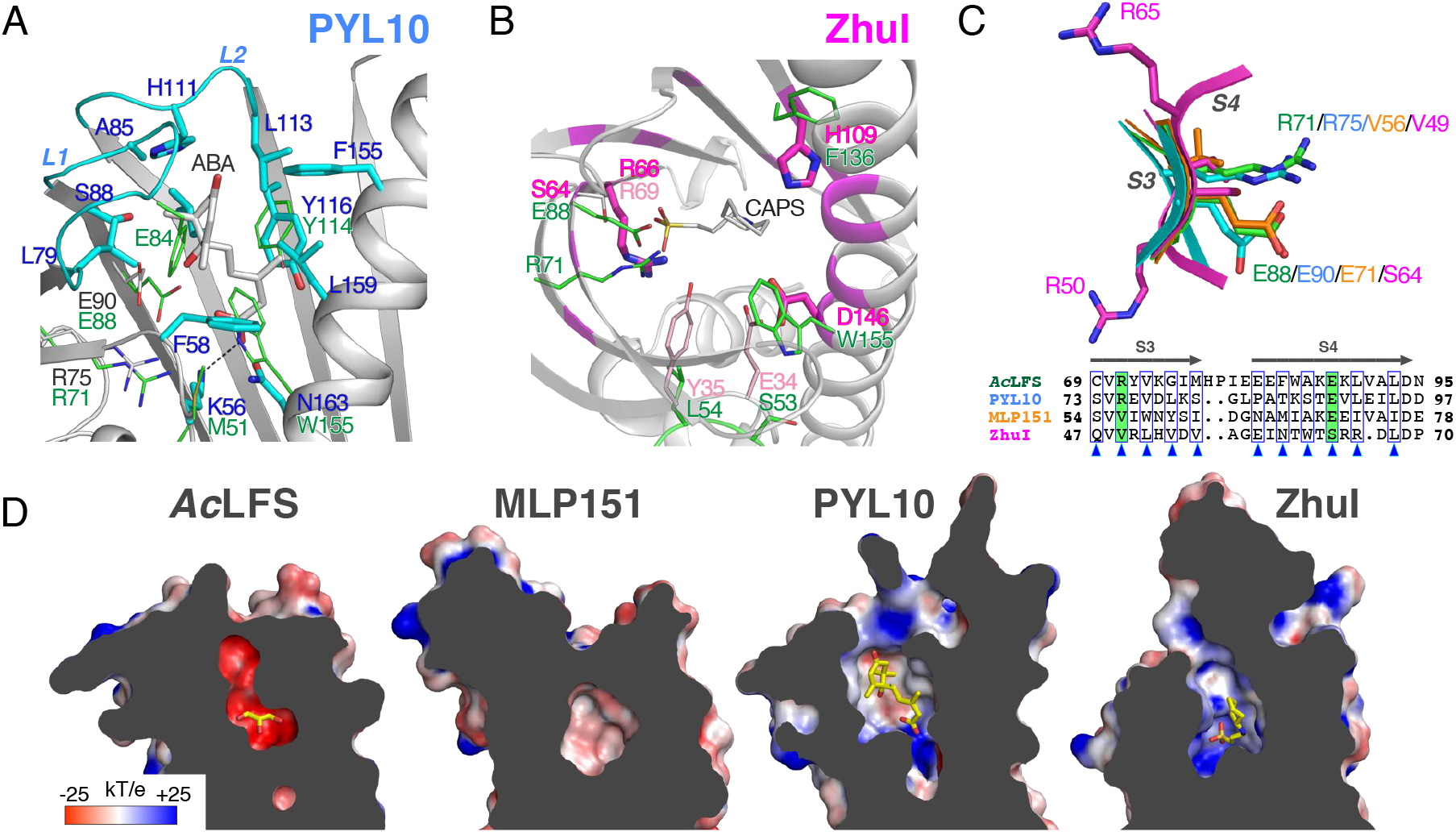
Comparisons with structural homologs. Close-up views of (*A*) PYL10-ABA (PDB 3r6p) and (*B*) ZhuI-CAPS (3tfz) complexes. ABA-contacting residues of PYL10 are colored cyan. Suggested primal catalytic residues of ZhuI and WhiE aro/cyc (PDB 3tvr) are colored magenta and pink. Corresponding residues of *Ac*LFS are overlaid as green sticks. (*C*) Superposition around the R71-E88 dyad motif of *Ac*LFS. Structure-based sequence alignment is depicted below. Arrowheads indicate residues whose side chains point inward. (*D*) Cutaway views of *Ac*LFS, MLP151 (PDB 4reh), PYL10, and ZhuI from left to right. Calculated electrostatic potentials are indicated on the surfaces following the order of blue to red.

Among the four proteins, conserved residues are mapped mainly on their structural frameworks rather than the active sites (Fig. S4). PYL10 harbors the ABA-binding site extending to peripheral L1 and L2 (Fig. 4*A*). ABA is recognized in the stretched conformation and a Floor residue K56 acts in orienting the carboxylate end (31). Three key residues of *Ac*LFS are fully conserved among PYL proteins, but R75, E90, and the hydroxyl group of Y116 do not directly contribute to the binding of ABA. Studies on mutagenesis and molecular docking have shown that the active site of ZhuI exists at a deeper position, around which a CAPS molecule is stabilized (34) (Fig. 4*D*). The substrate is thought to be placed in a compact conformation, which is ready for the nucleophilic attack from D146. R66 and H109 (Fig. 4*B*) are assumed to play an essential role in binding of the polyketide substrate (Fig. S3*A*). On the structures of WhiE aro/cyc and TcmN aro/cyc (41, 42), which are PKC isoforms possessing distinct regiospecificities toward dodeca- and decaketides, two of the deduced key residues needed for the corresponding catalytic reactions do not lie at the same positions: Y35 and E34 in WhiE aro/cyc instead of D146 and H109 in ZhuI (Fig. 4*B*). R66 is located on S3 and conserved among these PKCs (41). On S3 and S4, ZhuI has two other arginine residues, which are structurally not aligned on the R-E dyad motif of *Ac*LFS and PYL proteins (Fig. 4*C*), and the side chains of R50 and R65 are pointed away from the molecule by one-frame shifts. These findings indicate that the R-E dyad is a functional motif unique to LFS and not shared with the structural homolog proteins.

Although the key residues of SRPBCC proteins are totally divergent, the electrostatic landscapes in the pockets are comparable to the properties of respective acceptor molecules. In PYL10 and ZhuI, positive patches are distributed on nearly neutral intramolecular surfaces (Fig. 4*D*). They fit well with the carboxylate of ABA and negatively polarized acyl oxygen atoms of polyketide substrates. On the other hand, *Ac*LFS possesses an interior with comprehensively high negative potential, which would be effective at prohibiting the entry of anionic substances. In fact, a sulfate ion is visible only on the outer surface of *Ac*LFS (Fig. 3*B*). Sulfenic acid is regarded as being in protonation equilibrium (R-SOH ⇌ R-SO^−^ + H^+^) under physiological and aqueous conditions (13, 43). Therefore, the electronic filter is likely to have an advantage in isolating single 1-PSA in its protonated form, not 1-propene- 1-sulfenate.

### Proposed catalytic mechanism

Taking these findings together, a catalytic cycle of *Ac*LFS is proposed, as shown in Fig. 5*B*. The Wall apparatus first confines 1-PSA in a similar site of glycerol and 1,2-propanediol. The hydroxyl group of Y114 forms a hydrogen bond to the sulfenyl oxygen, and the adjacent hydrogen atom of the sulfenic acid orients toward the R71-E88 dyad, in which E88 constitutively rests in the dissociated form. The sulfenic acid is polarized and then deprotonated safely in the highly negative electrostatic potential inside the molecule, which is convenient for retention of a cation. The liberated proton forms a bond directly through the π electron pair of the 1-propenyl group, considering that there is no possible proton acceptor in the pocket. Indeed, the decrease in activity of E88D to the same level as its nonpolar variants indicates that *Ac*LFS would not exchange any electron pairs with the substrate (Fig. 2*D*). Y114 acts as not only a trap of 1-PSA but also an anchor of sulfenate anion, which is prone to leaving the site due to charge repulsion. Upon release of the product, Y114 no longer keeps PTSO in the pocket because of displacement around the S-linked oxygen, which arises from a valence change of the sulfur atom.

**Fig. 5.**
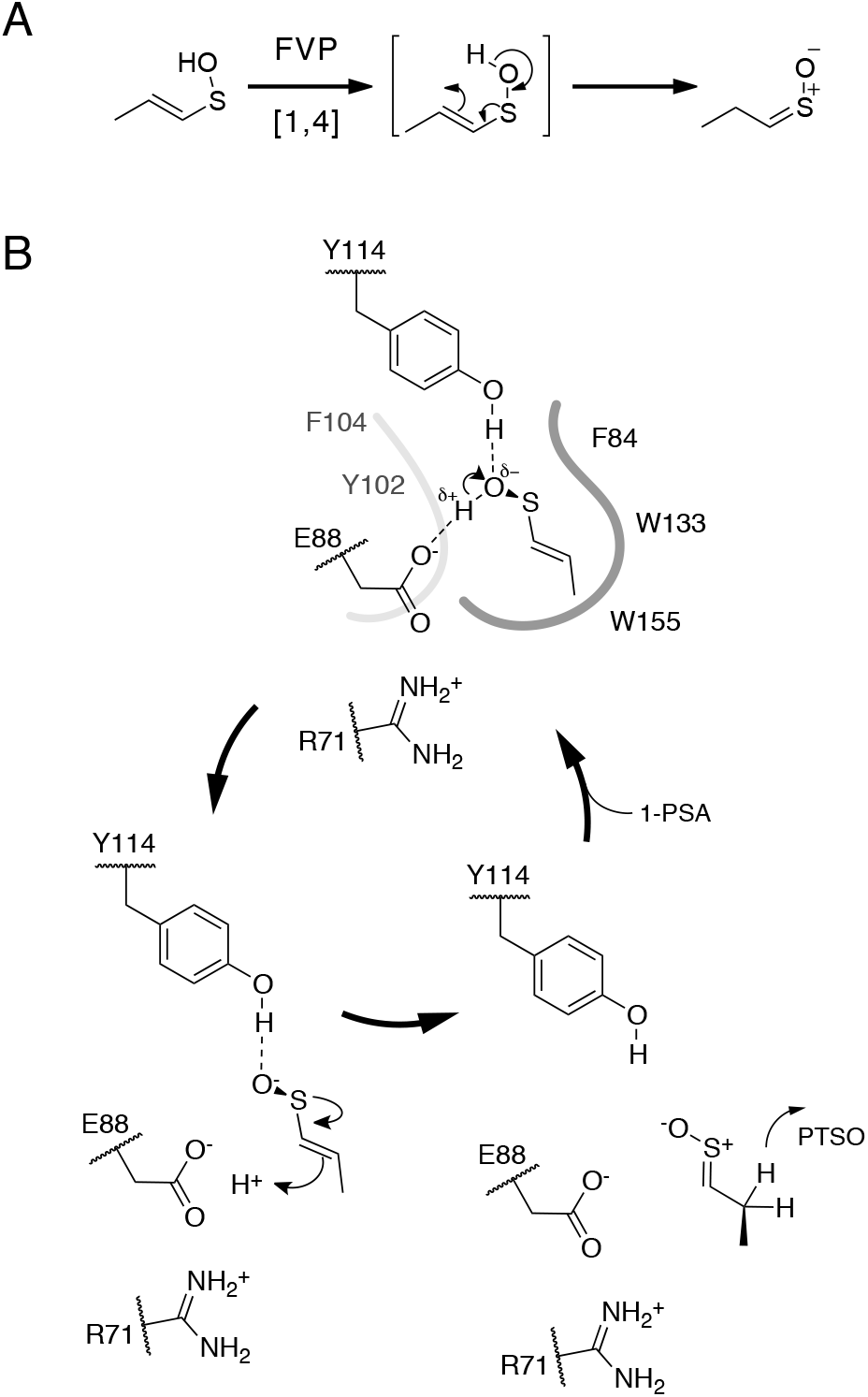
PTSO generation principles (*A*) under FVP condition and (*B*) with the assistance of *Ac*LFS.

The reaction scheme based on the intramolecular proton shuttle is consistent with a previous deuterium tracer study, which showed that 1-PSA is the proton source of PTSO (44). This scheme is apparently true in [1,4]-sigmatropic rearrangement, but the retro ene-reaction-like process occurs in a concerted fashion without yielding any intermediates (12) (Fig. 5*A*). *Ac*LFS provides a tailored compartment in order to increase the chances of correctly aligned overlap of binding orbitals, which is needed for the concerted reaction. Further, the compartment can precisely avoid self-condensation of 1-PSA monomers, involving a similar function to protection groups equipped in stable sulfenic acid compounds (45, 46). The roles in the catalysis are distinct from those of regular enzymes since the active site of *Ac*LFS is prepared for regulating the inherently labile nature of 1-PSA, rather than activating the substrates into respective transition states.

The present study provides a rationale for an RSS catalysis by revealing the three-dimensional structures of *Ac*LFS, which has a simple protein architecture. This work might not only decipher the principle of *Allium*-specific lachrymator generation, but also provide inspirations for multiple lines of work, particularly the synthesis of transient *S*-oxide compounds, which are not abundant and thus are rarely exploited.

## Materials and Methods

### Construction of plasmids

The sequences were amplified from a template plasmid harboring the complete open reading frame of *Ac*LFS (GenBank accession: AB089203). Fragments were digested with NdeI/XhoI and HindIII/BamHI, and then inserted into the corresponding sites of pET26b (Novagen) and pPAL7 (Bio-Rad) vectors to prepare the expression plasmids pET26b-*Ac*LFS and pPAL7-*Ac*LFS, respectively. Expression plasmids for *Ac*LFS mutants were prepared following the instructions of the PrimeStar mutagenesis kit (TaKaRa) using pET26b-*Ac*LFS as the template.

### Expression and purification of AcLFS

*Ac*LFS and its variants were expressed in *Escherichia coli* BL21AI cells (Life Technologies). Cells were cultured at 37°C in LB medium containing 50 mg/L kanamycin. After OD_600_ reached 0.6, *Ac*LFS was induced for 2 hours by adding a final concentration of 1 mM IPTG and 0.2% l-arabinose. Harvested cell pellets were suspended in solution A [50 mM Hepes-Na (pH7.5), 0.1 *μ*M PMSF] and disrupted by sonication (Sonifier 250D, Branson); then, insoluble particles were removed by centrifugation (20,000× g, 30 min). The supernatants were loaded onto HisTrap FF crude 5 mL (GE Healthcare) equilibrated with solution B [50 mM Hepes-Na (pH 7.5), 500 mM NaCl] and the eluates for solution C (solution B containing 500 mM imidazole) were pooled. The samples were loaded onto Superdex 200 10/300 GL (GE Healthcare) equilibrated with solution D [20 mM Hepes-Na (pH 7.5), 150 mM NaCl]. Peak fractions monitored by absorbance at 280 nm were collected and concentrated to around 20 mg/mL with the buffer exchanged by 20 mM Hepes-Na (pH 7.5) using Vivaspin turbo 10K MWCO (Sartorius).

For the production of selenomethionine (SeMet)-derivatized *Ac*LFS, pPAL7-*Ac*LFS was introduced into *E. coli* BL21 CodonPlus(DE3) RP-X cells. These cells were cultured in LeMaster medium containing 100 mg/L ampicillin and 35 mg/L chloramphenicol, and IPTG was added to the medium at a final concentration of 1 mM for induction. Extraction and purification were performed as per the protocols of non-derivatized *Ac*LFS, except for using a Profinity eXact column (Bio-Rad) instead of a HisTrap FF crude column, in accordance with the manufacturer’s instructions. In each step, purity was checked by SDS-PAGE and protein concentration was estimated using BCA protein assay kit (Thermo Fisher Scientific).

### Crystallography and structure determination

Crystallization was performed by the sitting-drop vapor diffusion method at 20°C. Crystals appeared within a week from the several batches of the conditions of the JCSG core I–IV and plus kits (Qiagen); then, the reservoir compositions were specifically optimized to the following condition: 2.0 M ammonium sulfate, 0.1 M MES-Na (pH 6.0). Each single crystals on the drops was picked out on CryoLoop (Hampton Research) and directly snapped in liquid nitrogen after cryo-protection, which was performed by transferring the crystal gradually to the reservoir solution containing 20% (v/v) glycerol or 16% D-trehalose. Immersion of the crystals in 1,2-propanediol [final 2% (v/v)] was performed together with the D-trehalose treatment.

Diffraction experiments were carried out at synchrotron facilities of KEK (Tsukuba, Japan), employing the mounting robot PAM (47) and client software UGUIS. The data acquisition scheme was basically set as follows: exposure of a 100 × 100 *μ*m^2^ beam for 1 sec per frame and the collection of 180 (360 for the SeMet dataset) frames with 1° oscillation steps. Diffraction images were indexed, integrated, and scaled using the HKL2000 program suite (HKL Research Inc.) or XDS (48).

Initial phases were determined by the single-wavelength anomalous dispersion (SAD) method using the SeMet dataset, practically by using the phenix.autosol module implemented in Phenix program suite (49). The resultant model was built mostly via phenix.autobuild and extended to the whole of the molecule manually, with viewing using the program Coot (50). Initial models corresponding to the other datasets were determined by molecular replacement, using the program Molrep (51) if needed. Models were refined iteratively using REFMAC5 in the CCP4 program suite (52, 53), and heteroatoms found were added according to *m*|*F*_o_|–*D*|*F*_c_| maps at the posterior rounds. The crystals originally belong to the *P*2_1_2_1_2_1_ space group, but are prone to being adapted as *P*4_1_ during pretreatment processes. The twin refinement option implemented in REFMAC5 was applied for the datasets indexed as the *P*4_1_ space group since nearly perfect twinning was found by phenix.xtriage. In addition, the occupancies of all of the atoms were set to 0.8 in the REFMAC5 calculations to reduce the phase bias-conducted lack of the resulting map densities around a region of one of the eight molecules in an asymmetric unit. The final model was verified by respective composite omit maps. Crystallographic and refinement statistics are listed in Table S1. Molecular graphics were prepared using PyMOL (Schrödinger LLC). Structure similarity searches were queued on Dali (54). Superposition and alignments of models were carried out on Coot. Electrostatic potentials were calculated using APBS (55) after charge state assignments with PROPKA (56). Coordinates and structure factors were deposited in the Protein Data Bank (PDB) with the accession codes of 5GTE (solute-free), 5GTF (with glycerol), and 5GTG (with 1,2-propanediol).

### Product assay

Liberated PTSO was isolated by reverse-phase chromatography at 25°C, as described previously (11). Briefly, a 3-min-reaction mixture was subjected to Pegasil ODS 4.5×250 (Senshu Kagaku) equilibrated with 0.01% trifluoroacetic acid in 30% methanol solution (pH~3.3) and then developed for 12 min with a flow rate of 0.6 mL/min. Absorbance at 254 nm was monitored using an Alliance instrument (Waters). A total batch of 25 *μ*L initially consisted of 5 *μ*L of 0.01 mg/mL *Ac*LFS, 5 *μ*L of respective inhibitor in 5-fold concentrate and 10 *μ*L of garlic alliinase (250 U/mL), then 5 *μ*L of 20 mg/mL *trans*-1-PRENCSO was added to start the reaction. Cysteine thiols on *Ac*LFS were inactivated by adding iodoacetamide at a final concentration of 1 mM in 0.01 mg/mL *Ac*LFS and incubating the mixture at 4°C for several hours in the dark. Chemical reagents were purchased from Wako Pure Chemicals and Tokyo Chemical Industry.

## Author contributions

S.F. conceived the research and T.A. designed experiments; J.T., Y.S., and T.A. performed research; N.M., M.K., M.A, T.K., N.T., and S.I. supplied *Ac*LFS gene and materials; All the authors discussed data; T.A. wrote the paper with assistance from all other authors.

The authors declare no conflict of interest.

## Acknowledgments

The authors would thank to the beamline scientists of KEK and SPring-8. Also thank to Drs. Yoshitaka Moriwaki and Kentaro Shimizu for discussion and preliminary simulations. This work was supported by JSPS KAKENHI Grant Numbers JP26660289 (to T.A.), JP15H02443, and JP26660083 (to S.F.).

## Supplemental information

**Table S1.**
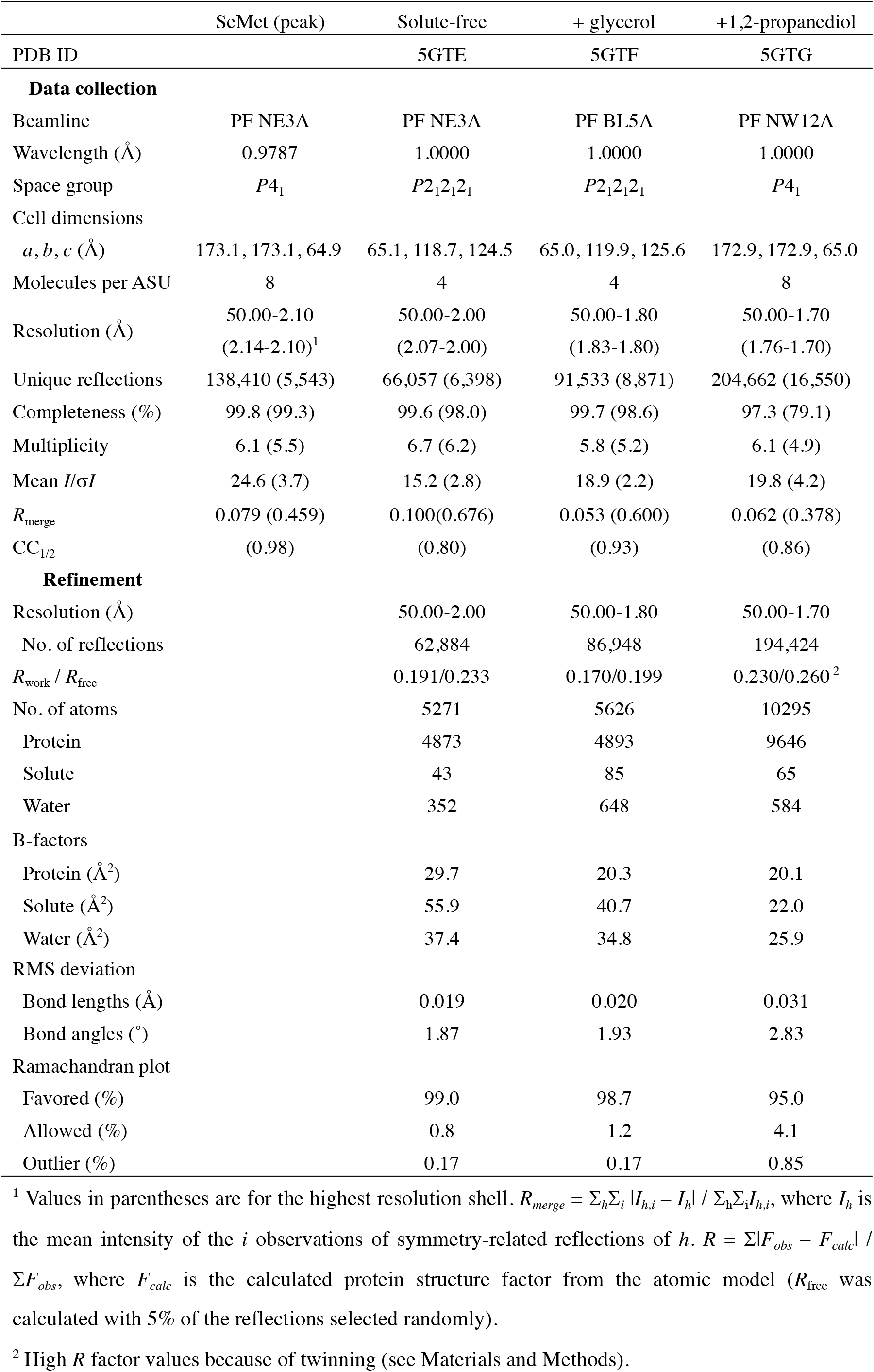
Data collection and refinement statistics

**Table S2.** Proteins of similar structure to *Ac*LFS^1^

| Name | Source | Z<sub>max</sub> | Identity <sup>2</sup> | Similarity <sup>2</sup> | Domain family <sup>3</sup> | PDB ID <sup>4</sup> |
| --- | --- | --- | --- | --- | --- | --- |
| PYL10 | *Arabidopsis thaliana* | 19.0 | 16.7%<br>(31/186) | 32.3%<br>(60/186) | PYR_PYL_RCAR_like<br>(cd07821) | 3R6P |
| PYL9 | *A. thaliana* | 17.3 | 17.3%<br>(33/191) | 30.4%<br>(58/191) | PYR_PYL_RCAR_like | 3OQU |
| MLP151 | *Panax ginseng* | 17.2 | 18.8%<br>(32/170) | 34.1%<br>(58/170) | Bet_v_1_like<br>(cd07816/smart01037) | 4REH |
| PYL2 | *A. thaliana* | 17.1 | 16.7%<br>(32/192) | 32.3%<br>(62/192) | PYR_PYL_RCAR_like | 3KB0 |
| PYL1 | *A. thaliana* | 16.9 | 16.8%<br>(30/179) | 31.8%<br>(57/179) | PYR_PYL_RCAR_like | 3JRS |
| ZhuI | *Streptomyces*<br>sp. R1128 | 16.6 | 12.1%<br>(21/174) | 24.7%<br>(43/174) | OtcD1_ARO-CYC_like<br>(cd08861) | 3TFZ |
| PYL3 | *A. thaliana* | 16.5 | 17.2%<br>(36/209) | 31.6%<br>(66/209) | PYR_PYL_RCAR_like | 4DSC |
| WhiE aro/cyc | *S. coelicolor*<br>A3(2) | 15.7 | 14.1%<br>(25/177) | 27.7%<br>(49/177) | OtcD1_ARO-CYC_like | 3TVQ |

<sup>1</sup> Sorted by a descending order of the maximum Z-scores of DaliLite v. 3.
<sup>2</sup> Taken from the output of EMBOSS Stretcher.
<sup>3</sup> Acquired from Conserved Domain Database (CDD).
<sup>4</sup> Representative entry (ligand complex if available) is shown.

### Figure legends

**Figure S1.**
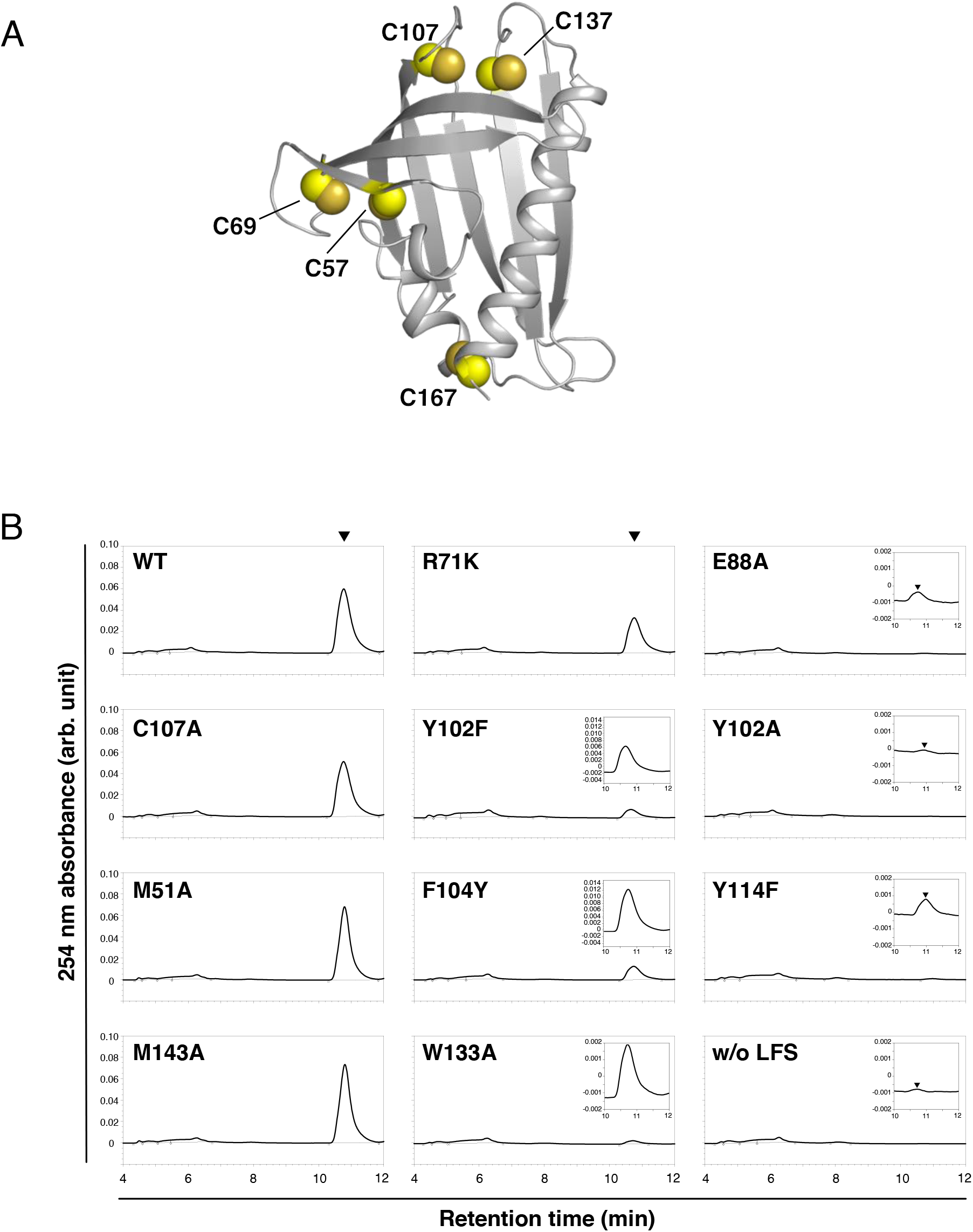
(*A*) Distribution of cysteine residues on *Ac*LFS structure. (*B*) chromatograms of the selected measurements of reaction products. Arrowheads indicate the retention times of PTSO.

**Figure S2.**
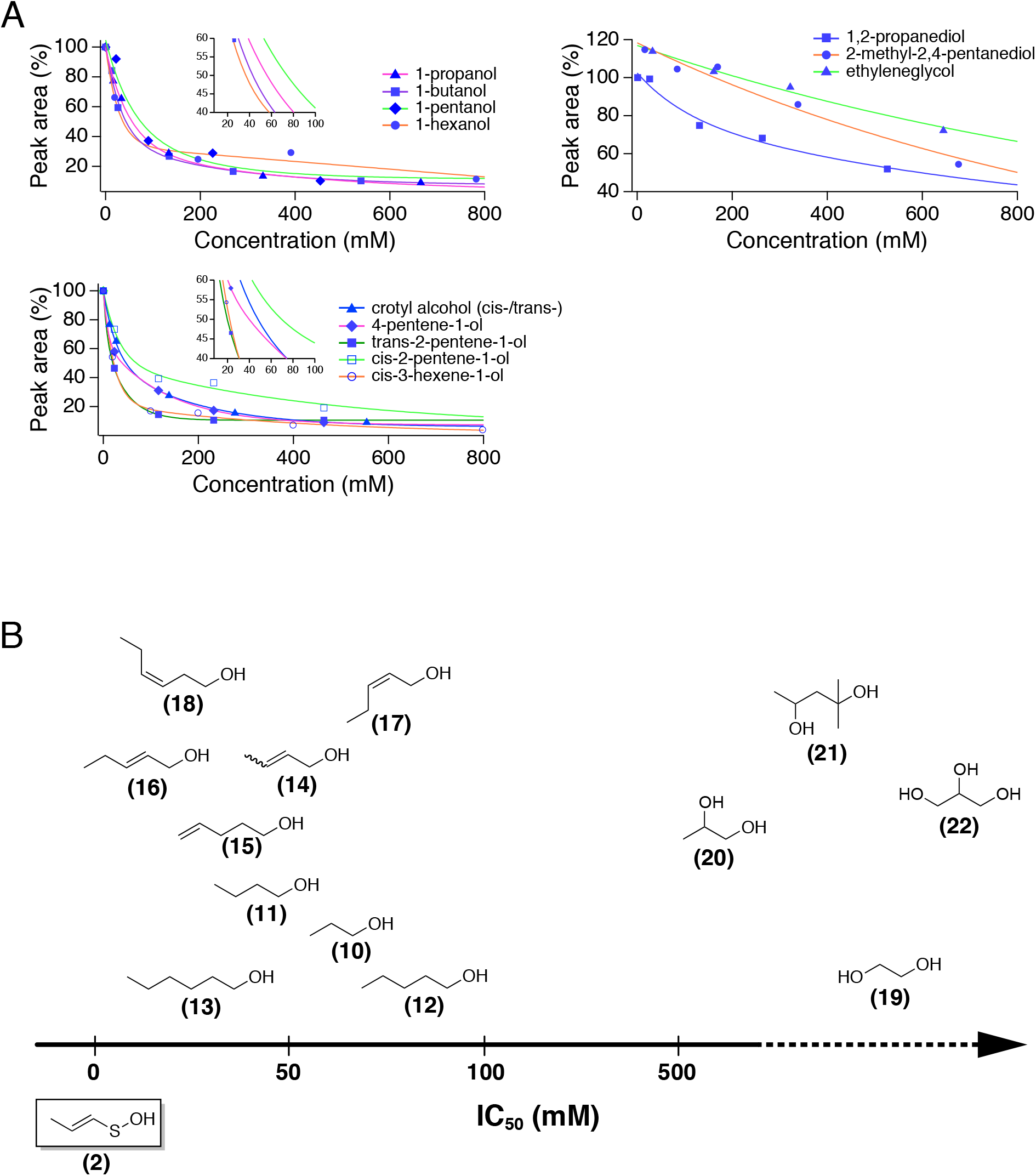
Inhibitions by organic hydroxide compounds. (*A*) Inhibition curves in the presence of normal alcohols (top), unsaturated alcohols (bottom) and diols (right). (*B*) Array according to the estimated IC_50_ values of solute molecules tested in this study. Structural formulae stand for 1-propanol (**10**), 1-butanol (**11**), 1-pentanol (**12**), 1-hexanol (**13**), crotyl alcohol (**14**), 4-pentene-1-ol (**15**), *trans*-2-pentene-1-ol (**16**), *cis*-2-pentene-1-ol (**17**), *cis*-2-hexene-1-ol (**18**), ethylene glycol (**19**), 1,2-propanediol (**20**), 2-methyl-2,4-pentanediol (**21**) and glycerol (**22**).

**Figure S3.**
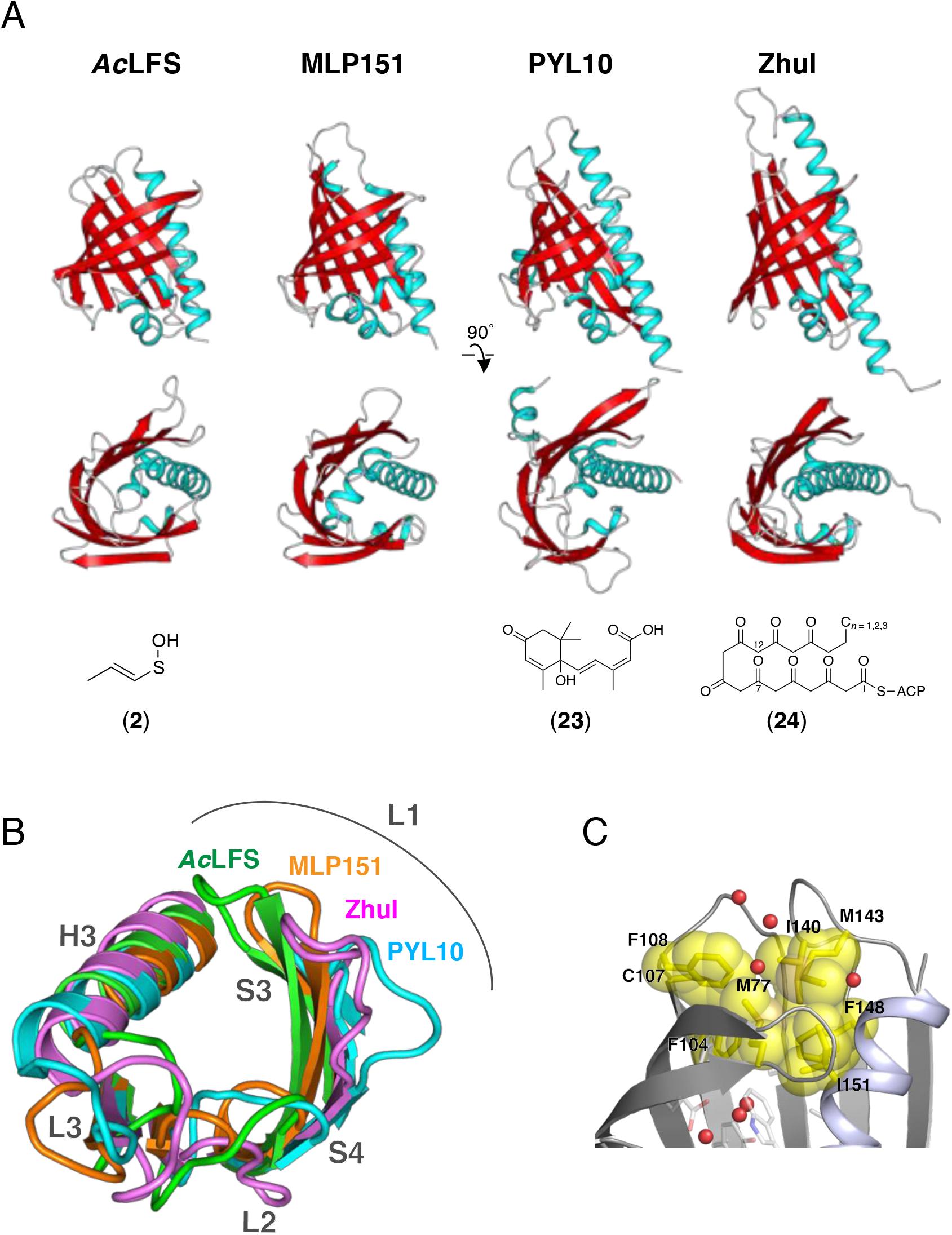
(*A*) Comparison of structurally relative helix-grip fold proteins. Primary substrate or ligand of each protein is displayed at the bottom. Structural formulae stand for abscisic acid (**23**) and octaketide–acyl carrier protein conjugate (**24**). (*B*) Superposition around the entrance. Cartoon representations of *Ac*LFS (green), MLP15 (orange), PYL10 (cyan) and ZhuI (magenta). Superposed view from Floor to Gate of the pocket. (*C*) Cluster of hydrophobic residues on Gate layer. Spheres are drawn as respective van der Waals radii of carbon (yellow) and sulfur (orange).

**Figure S4.**
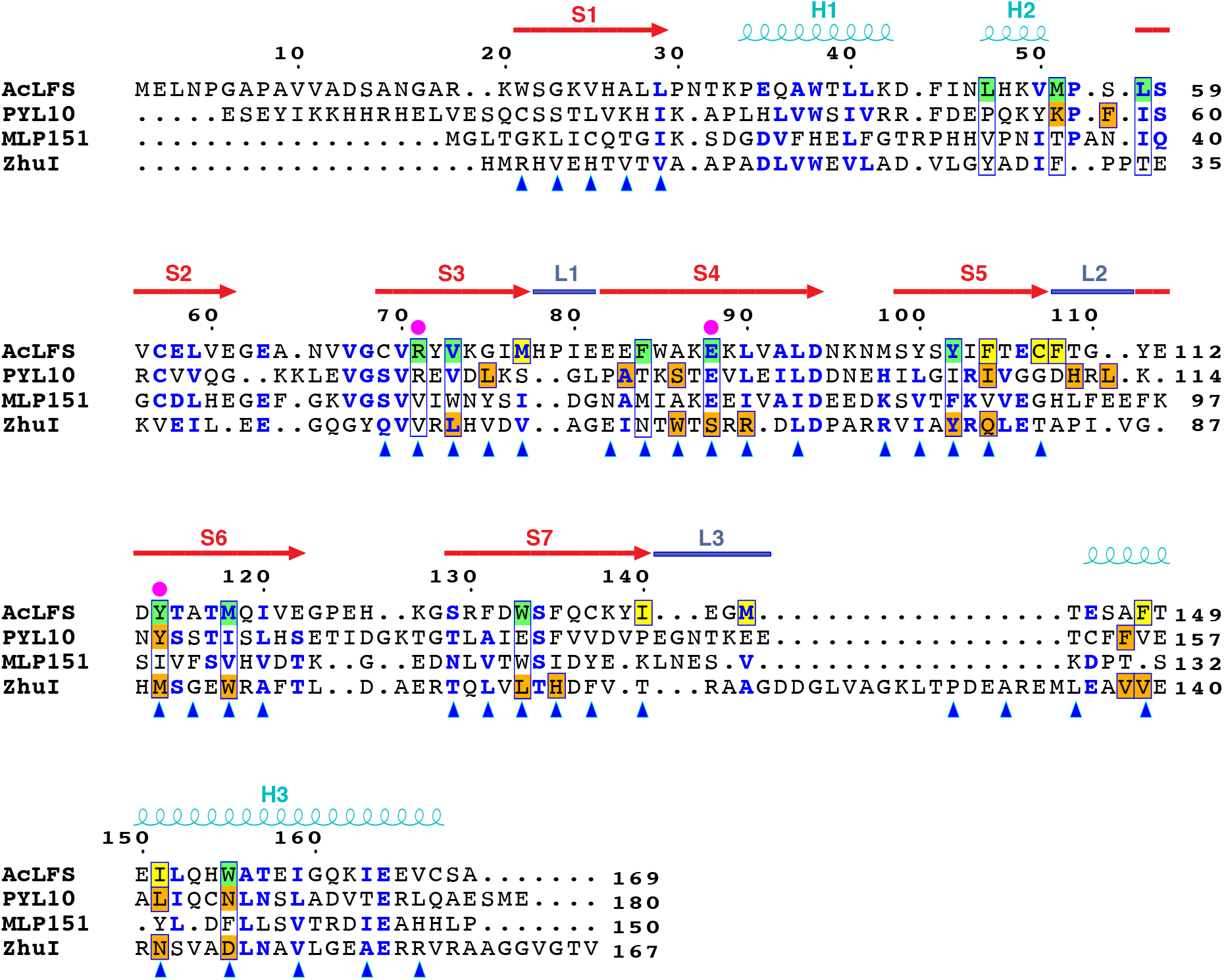
Structure-based amino acid sequence alignment of *Ac*LFS and structural homologs. Conserved residues represent blue characters. Residues are highlighted for Wall and Floor (green), Gate (yellow), direct association with ABA (orange in PYL10) and CAPS (orange in ZhuI). The key residues of *Ac*LFS are marked with magenta dots and secondary structure elements drawn upside are for *Ac*LFS. Arrowheads indicate residues pointing inward.

